# Using Touchscreen Equipped Operant Chambers to Study Comparative Cognition. Benefits, Limitations, and Advice

**DOI:** 10.1101/2020.10.03.324814

**Authors:** BM Seitz, KB McCune, M MacPherson, L Bergeron, AP Blaisdell, CJ Logan

## Abstract

Operant chambers are small enclosures used to test animal behavior and cognition. While traditionally reliant on simple technologies for presenting stimuli (e.g., lights and sounds) and recording responses made to basic manipulanda (e.g., levers and buttons), an increasing number of researchers are beginning to use Touchscreen-equipped Operant Chambers (TOCs). These TOCs have obvious advantages, namely by allowing researchers to present a near infinite number of stimuli as well as increased flexibility in the types of responses that can be made and recorded. Here, we trained wild-caught adult and juvenile great-tailed grackles (*Quiscalus mexicanus*) to complete experiments using a TOC. We have learned much from these efforts, and outline the advantages and disadvantages of these two approaches. We report data from our training sessions and discuss important modifications we made to facilitate animal engagement and participation in various tasks. Finally, we provide a “training guide” for creating experiments using PsychoPy, a free and open-source software that we have found to be incredibly useful during these endeavors. This article, therefore, should serve as a useful resource to those interested in switching to or maintaining a TOC, or who similarly wish to use a TOC to test the cognitive abilities of non-model species or wild-caught individuals.

---

A number of scientific disciplines including behavioral neuroscience, experimental psychology, ethology, and ecology, aim to understand the cognitive abilities of animals. Given the so-called *black box paradox* of studying cognition, where behavioral measures are used to infer cognitive abilities, a number of technologies have been developed that measure behavioral responses to a variety of stimulus inputs. However, these technologies have been primarily used with captive-reared model species, thus limiting our ability to generalize results to naturalistic conditions (due to the intentional or unintentional artificial selection that is often involved; Niemela & Dingemanse, 2014) and make cross-species generalizations about cognitive abilities. Perhaps the most influential technology developed to study animal cognition is the “operant chamber” (also referred to as the *Skinner Box*). While many variations exist, an operant chamber (OC) typically consists of a small enclosure with one or several manipulanda (e.g. levers, chains) and responses made to them are recorded by a computer. Critical to the OC is also the ability to present a variety of stimuli (e.g., auditory and visual cues) as well as biologically significant outcomes (e.g., food or shock) that are often contingent on the animal’s interactions with the manipulanda.

As technologies advance, many researchers have transitioned to using touchscreen-equipped operant chambers (TOCs) because they confer a number of advantages (see below). TOCs are now commonplace in studies utilizing more traditional laboratory animals such as rodents, primates, and pigeons. TOCs have also been used to study non-model species such as bears, dogs, and tortoises in captivity (Vonk & Beran 2012; Perdue 2016; Zeagler et al. 2016; Mueller-Paul et al. 2014). However, rarely has the use of this method been described in wild-caught individuals from non-model species (but see Chow et al., 2017; Guigueno, MacDougall-Shackleton, & Sherry, 2015). There is a paucity of information about how to train such individuals to use a TOC, potentially because OCs and TOCs were tailored to the behavior and characteristics of model species, which could explain the difficulty in training non-model species and the increased variability in training success and performance. This distinction is relevant to the ability to generalize TOC procedures across groups because captive-reared animals often have different ontogenetic experiences that affect their motivation to interact with, and performance on, behavioral tasks compared to wild conspecifics (McCune, Jablonski, Lee, & Ha, 2019; Vardi et al., 2020). We embarked on an investigation using TOCs in wild-caught adult great-tailed grackles (*Quiscalus mexicanus*; hereafter grackles) that were temporarily held in aviaries. We discovered that extensive modifications to procedures used with laboratory pigeons were required to train grackles to use such an apparatus. Our aim here is to briefly describe the advantages and disadvantages of TOCs for animal cognitive testing, report data summarizing our efforts to train several grackles, and finally, to provide a “training guide” to facilitate the use of TOCs in non-model species while using open source software.

## Advantages and Disadvantages of Touchscreen-Equipped Operant Chambers (TOCs)

### Advantages of TOCs

The rise of TOCs in behavioral research is likely due to a multitude of factors. First, touchscreen technology has advanced greatly in the past several decades, and the application of touchscreens in everyday devices (e.g., mobile phones, tablets, ATMs) has become commonplace. This makes converting such technologies for research purposes particularly appealing. Additionally, there are a number of advantages associated with using TOCs compared to traditional OCs or open field tasks. Notably, TOCs allow for the presentation of virtually any combination of visual and/or auditory stimuli (Kangas & Bergman, 2017; Morrison & Brown, 1990).This feature can be advantageous for studying animals over a prolonged period of their lifespan because typical OCs only have a limited number of stimuli that can be presented (e.g., various wavelengths of light). This can be problematic if animals generalize stimuli from task to task, but this is mitigated when using a TOC due to the near infinite visual stimulus types that can be presented. Further, TOCs allow novel stimulus types such as video to be presented to animals. This facilitates the investigation of a much wider range of phenomena compared to what can typically be studied using a traditional OC (e.g., social learning, predator/prey learning, motion processing). Note, these specific advantages can be accomplished simply by adding a computer monitor to an existing experimental setup and do not necessitate the complete TOC setup (for examples, see Shragrai et al., 2017; Wood & Wood, 2016), and researchers must consider the perceptual capabilities (if known; if not, then speculation could help) of their model organism before employing video stimuli (D’Eath, 1998; Lea & Dittrich, 2000). Additionally, while the number of visual stimuli that can be presented is drastically increased, simple TOCs offer no advantage in terms of other, diverse stimulus characteristics (e.g., scent, taste, texture, temperature).

Whereas OCs typically have a limited number of spatially-fixed response options (e.g., lever pressing or chain pulling), TOCs again afford a much wider range of possible response locations (Morrison & Brown, 1990). This is beneficial not only when testing the same animal on multiple tasks but also for adjusting potential side biases and other response biases which are commonplace in operant conditioning procedures (Kangas & Bergman, 2017). TOCs record not only that a response has been made, but also, precisely when in the trial and where on the screen the response was made, unlike traditional manipulanda found in OCs (Stahlman, Young, & Blaisdell, 2010). As such, TOCs are particularly useful for investigations of spatial learning and memory processes (Sawa, Leising, & Blaisdell, 2005; Wolf, Urbano, Ruprecht, & Leising, 2014). Given the TOC recording abilities and flexibility of stimulus presentation, they can be used to study phenomena potentially governed by more complex cognitive processes—or at least, phenomena that involve more complex procedures (e.g., Katz & Wright, 2006)—while traditional OCs are generally limited to tasks comprising of Pavlovian (stimulus-outcome associations) and instrumental (response-outcome or stimulus-response associations) processes—which involve simple procedural designs.

Removing the experimenter from the subject’s periphery also reduces potential experimenter-induced bias (e.g., the experimenter’s unconscious body language might unknowingly cue the subject to make the correct choice). While this is also a feature of typical OCs, TOCs can be remotely controlled by an experimenter outside the periphery of the subject (see section “*The Touchscreen-equipped Operant Chamber*” for more on this), allowing for even more control over the function of the program and protecting against instances when the individual is not attending to or not motivated to perform the task. In this manner, TOCs offer convenience to researchers because sessions are easy to initiate, and trials can be programmed to begin automatically or manually without disrupting the subject. Responses are measured reliably, and subjects can complete numerous consecutive trials depending on the experimental parameters.

TOCs also allow for experimental procedures (e.g., protocols, code, methods) to be more easily shared between researchers because the software programs for developing and running TOC experiments can be shared and modified, though we, and others (e.g. Dumont, Salewski, & Beraldo, 2020), argue there is still much room for improvement on this front. The ease of sharing programs and data sets among research groups around the world, and the flexibility in stimuli and testing paradigms should facilitate the inclusion of novel species trained with TOCs as well as cross-species comparisons. As an example, Seitz, Stolyarova, & Blaisdell (2020) tested pigeon and human performance on a nearly identical TOC procedure. The only difference between the two experiments was that pigeons received a grain reward and humans accumulated points that could later be traded for candy. A number of species have already been successfully trained using TOCs, which points to the flexible and intuitive nature of touchscreen tasks (Kangas & Bergman, 2017; Schmitt, 2018). In fact, there is evidence that touchscreens can be more effective in training animals on discrimination-based tasks than traditional OC methods, presumably because touchscreens allow the animal to interact directly with the stimulus to obtain some immediate outcome (i.e., touch the stimulus to make a choice) instead of responding to a lever that is spatially separated from the stimulus (Bussey et al., 2008; Cook, Geller, Zhang, & Gowda, 2004).

### Disadvantages of TOCs

Despite the advantages associated with TOCs, their use in studying cognitive abilities in animals poses a number of new and unique challenges that should be considered by researchers unfamiliar with this method. The most obvious barrier to adopting TOCs is that they require researchers to possess some level of programming skills. This can be incredibly difficult for researchers whose backgrounds are far removed from computer science. Some researchers have circumvented this issue by hiring programmers to build their programs, but this can be a costly (around ∼$1,000 per week) and inefficient solution. Moreover, the existence of multiple different programming languages, some of which are not free to use (e.g., E-Prime or MATLAB), makes it very difficult for experimenters to share experimental paradigms with each other. This reduces the possibility of collaboration as well as the potential to replicate the research of others, both of which hinder scientific progress. We later discuss the benefits of using Psychopy (Psychopy.org), an open-source programming software for creating experiments for TOCs as a solution to these challenges.

Another challenge we’ve encountered using TOCs are accidental contacts made to the touchscreen which are then measured as responses. This often occurs when the bird’s breast or tail makes contact with the screen. This is a potentially serious issue because unintentional actions that lead to rewards or punishment can drastically affect learning processes and behavior (Skinner, 1948). Fortunately, these programs can be easily formatted such that only responses made to specific stimuli (and not the entire screen) are recorded and/or result in some specific outcome. That said, accidental contact to stimuli does occasionally occur, which suggests that coupling this technology with a video recording device can be particularly helpful. Related to this, at least with avian species in our specific setup (see details below), not all pecks to the TOC appear to be registered by the infrared system. This does not have much impact during the later stages of training, because birds will often make multiple pecks to the same stimulus, but it can be a challenge during initial training. Further, when a response is accidentally made or missed, there are limited ways for correcting for this in real time (but see below for our suggested technique of coupling the experimental program with a livestream video call to remote control the operating computer). Researchers interested in building their own TOCs may wish to consult engineers to ensure their touchscreen equipment is sensitive to the types of responses made by their respective species of study.

## Lessons from Teaching Wild-Caught Animals to Use Touchscreens

For the past two years, we (Logan et al. 2019a, b, Blaisdell et al. 2019) have been investigating the behavior of wild-caught grackles, using a number of different experimental procedures, many of which involved using a TOC. We encountered several unexpected hurdles in training grackles to use the TOC, which could be due to species differences and/or because they were wild-caught rather than captive-bred. Consequently, we gained valuable insights that are likely useful for other researchers interested in a similar approach. These insights cluster around two major themes: 1. Managing challenges that arise from training wild-caught adult birds to operate a TOC compared to traditional laboratory animals like pigeons; and 2. The benefit of using PsychoPy software to create behavioral tasks for TOCs. We discuss our process of navigating these themes below, we use our TOC training data to posit relationships with training performance and the number of prior non-TOC experiments and TOC experiments completed, and finally detail our process for using PsychoPy software to design behavioral tasks on TOCs.

TOC testing opened up new avenues for the types of comparative cognition experiments we were able to conduct with temporarily-held wild birds, but which required several modifications to allow the successful testing of grackles. Our detailed grackle TOC training protocol can be found here: https://docs.google.com/document/d/18HJzHVk-iFUwpBYRQoxVy1J6izFFr1_2joS9CRkyNSM/edit?usp=sharing. By incorporating TOC tests into our test battery, we were able to test individuals with less interference from human experimenters (making testing faster), conduct tests that were difficult to implement without a TOC (e.g., Go/No-Go and causal cognition tests), and validate the ecological relevance of previous TOC experiments that used individuals from model species that have been captive for generations and thus might lack the behavioral responses that one would see in wild individuals that are subject to natural selection (Morand-Ferron & Quinn 2016, Mery 2013, Kamil 1994). Additionally, if we can effectively modify the apparatus and training, it could allow us to bring the TOC to the field and eliminate the need for these experiments to occur in captivity (e.g., for wild vs. captive reversal learning using operant boxes, see Cauchoix et al 2017; for examples of automated feeders used in the wild, see Morand-Ferron et al. 2015 and Aplin 2018).

### The Touchscreen-equipped Operant Chamber

Our TOC consisted of a color LCD monitor (NEC MultiSync LCD1550M) measuring 23.2 cm x 30.5 cm that sat behind an infrared touchscreen (Carroll Touch, Elotouch Systems, Fremont, CA). A food hopper was located below the monitor with an access hole situated parallel to the floor. The hopper was raised and lowered using a robotic arm (Pololu Maestro 12 Channel USB Servo, Pololu Robotics, Las Vegas, NV) and, when in the raised position, the food inside the hopper was accessible. While a TOC set-up is typically enclosed by an opaque chamber, this was removed for the grackles to allow the birds to engage with the apparatus at will and thus not rely on researchers actively placing the bird in the OC for participation (see figure 1 for a picture of our setup during hopper training). One major piece of advice we offer is installing software (e.g., Teamviewer or Microsoft Remote Desktop) that allows one to remotely control the computer being used to operate the TOC. This provides a solution for the issue of a response not being detected by the touchscreen because the experimenter can use a different computer to register the response made by the animal on the TOC—either by observing from a distance or over live video feed. For example, by simply installing the TOC with a webcam and initiating a Skype (www.skype.com) call with the remotely-controlled computer, we are able to observe the bird’s behavior in real time and intervene if necessary. For a detailed description of the grackle training procedures including a discussion of techniques and programs that did/did not work, see *S1*.

**Figure 1.**
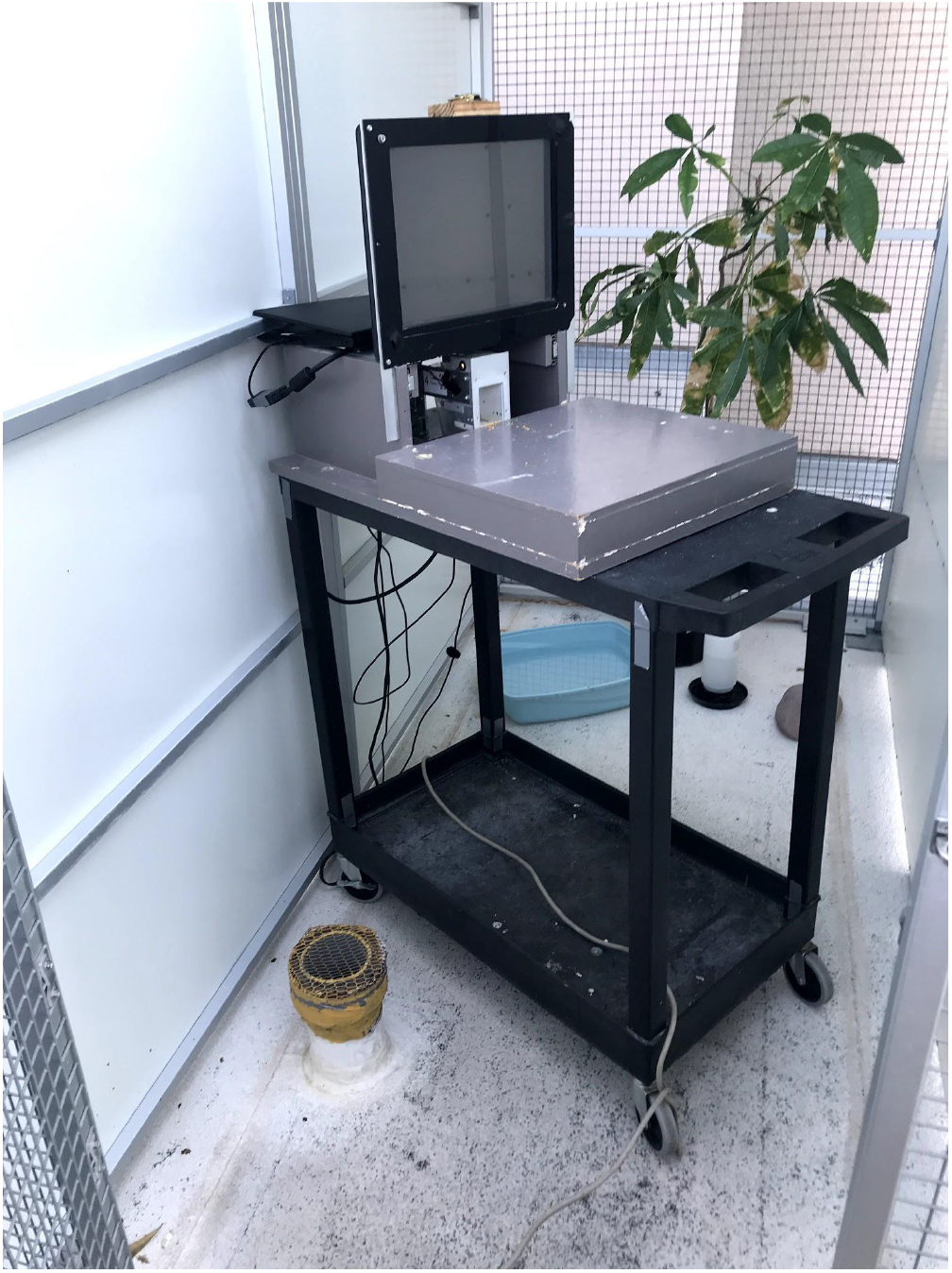
Touchscreen-equipped operant chamber basic set up during food hopper training. The computer monitor sits behind a touchscreen panel, and food is delivered via a food hopper that is moved by a robotic arm. The programs are run on a laptop located at the back of the touchscreen, and the laptop can be remotely controlled from outside the aviary using the experimenter’s laptop. Unlike most TOCs, our setup is not an enclosed chamber.

#### General Touchscreen-equipped Operant Chamber Training Rules

Throughout the training process and during testing, it was crucial to make the TOC apparatus available to the bird only when we wanted the bird to participate in a training trial. During daily testing and training, if the bird did not interact with the TOC within 5 minutes of it being placed inside their aviary, the TOC was removed and we tried again later. This resulted in the birds learning that if they wanted to participate, they must do so right away. Additionally, when they did choose to participate, we would stop the session and remove the TOC if they got distracted by interacting with different parts of the TOC or other objects in their aviary, or lost motivation and did not return to the TOC within 5 minutes. This resulted in the birds learning that if the TOC is available to them, they must stay engaged with it until completing the test. Consequently, testing sessions were faster and more concentrated than if experimenters had waited for birds to participate.

### Questions our limited training data can begin to answer (post-hoc)

The data we collected during TOC training allowed us to begin to qualitatively investigate a few *post-hoc* insights (Table 1). Such insights might be useful for programs that already run TOC experiments to determine whether there are ways to improve efficiency (e.g., what leads to faster training times) and how to decide which individuals to select to participate in these experiments. These insights could also help a researcher determine whether it would be feasible to implement a TOC experiment given the amount of time that is necessary for training. To answer the below questions, we examined the data in Table 1 for those birds who completed enough of the training to determine whether they were a fast (<13 days to pass training) or slow learner (13+ days to pass training) (n=11 grackles).

**Table 1.**
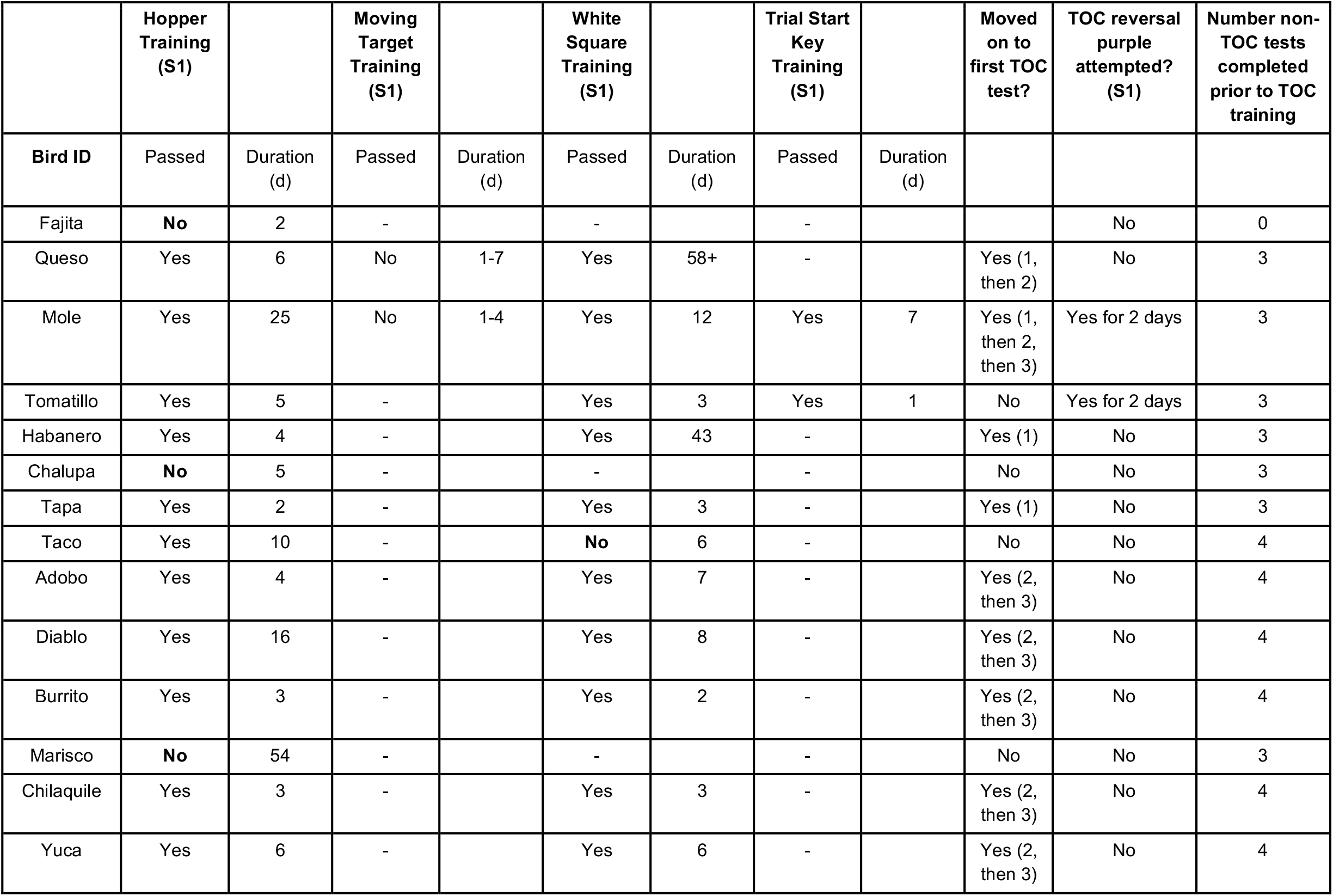
Summary data for grackles that went through TOC training. Duration=number of days. In ‘Moved on to first TOC test’: 1=reversal TOC experiment, 2=Go/No-Go TOC experiment, 3=causal cognition TOC experiment. Some birds did not pass training because they had to be released back to the wild (Fajita, Tomatillo, Chalupa, and Taco); Queso had a medical procedure in the middle of white square training, which extended his training period because we waited for him to recover and then to become motivated to participate again; Marisco was not close to passing hopper training after 54 days, so we stopped trying. Note: the maximum possible number of TOC experiments was three from birds Fajita through Tapa, and two thereafter. The maximum possible number of non-TOC tests completed prior to TOC training was three from birds Fajita through Tapa, and four thereafter.

|  | Hopper Training (S1) |  | Moving Target Training (S1) |  | White Square Training (S1) |  | Trial Start Key Training (S1) |  | Moved on to first TOC test? | TOC reversal purple attempted? (S1) | Number non-TOC tests completed prior to TOC training |
| --- | --- | --- | --- | --- | --- | --- | --- | --- | --- | --- | --- |
| Bird ID | Passed | Duration (d) | Passed | Duration (d) | Passed | Duration (d) | Passed | Duration (d) |  |  |  |
| Fajita | **No** | 2 | - |  | - |  | - |  |  | No | 0 |
| Queso | Yes | 6 | No | 1-7 | Yes | 58+ | - |  | Yes (1, then 2) | No | 3 |
| Mole | Yes | 25 | No | 1-4 | Yes | 12 | Yes | 7 | Yes (1, then 2, then 3) | Yes for 2 days | 3 |
| Tomatillo | Yes | 5 | - |  | Yes | 3 | Yes | 1 | No | Yes for 2 days | 3 |
| Habanero | Yes | 4 | - |  | Yes | 43 | - |  | Yes (1) | No | 3 |
| Chalupa | **No** | 5 | - |  | - |  | - |  | No | No | 3 |
| Tapa | Yes | 2 | - |  | Yes | 3 | - |  | Yes (1) | No | 3 |
| Taco | Yes | 10 | - |  | **No** | 6 | - |  | No | No | 4 |
| Adobo | Yes | 4 | - |  | Yes | 7 | - |  | Yes (2, then 3) | No | 4 |
| Diablo | Yes | 16 | - |  | Yes | 8 | - |  | Yes (2, then 3) | No | 4 |
| Burrito | Yes | 3 | - |  | Yes | 2 | - |  | Yes (2, then 3) | No | 4 |
| Marisco | **No** | 54 | - |  | - |  | - |  | No | No | 3 |
| Chilaquile | Yes | 3 | - |  | Yes | 3 | - |  | Yes (2, then 3) | No | 4 |
| Yuca | Yes | 6 | - |  | Yes | 6 | - |  | Yes (2, then 3) | No | 4 |

Question 1: Were those grackles that completed more experiments prior to beginning TOC training more likely to **complete TOC training faster**? If the answer to this question (or question 2) is yes, this is potentially because more test-taking experience leads to individuals becoming better test takers, and/or more habituated to the aviary testing environment. In these cases, experimenters should put the TOC tests at the end of a test battery that involves both TOC and traditional procedures, or test individuals who already have extensive prior testing experience. Answer 1: The number of previously completed experiments (three or four) was likely not related to TOC training duration (Table 2). All grackles, except Marisco, completed the maximum number of non-TOC tests possible before beginning TOC training and their training durations varied, sometimes substantially. Thus, participating in more experiments before beginning TOC training did not appear to result in faster, and thus more efficient, training.

**Table 2.**
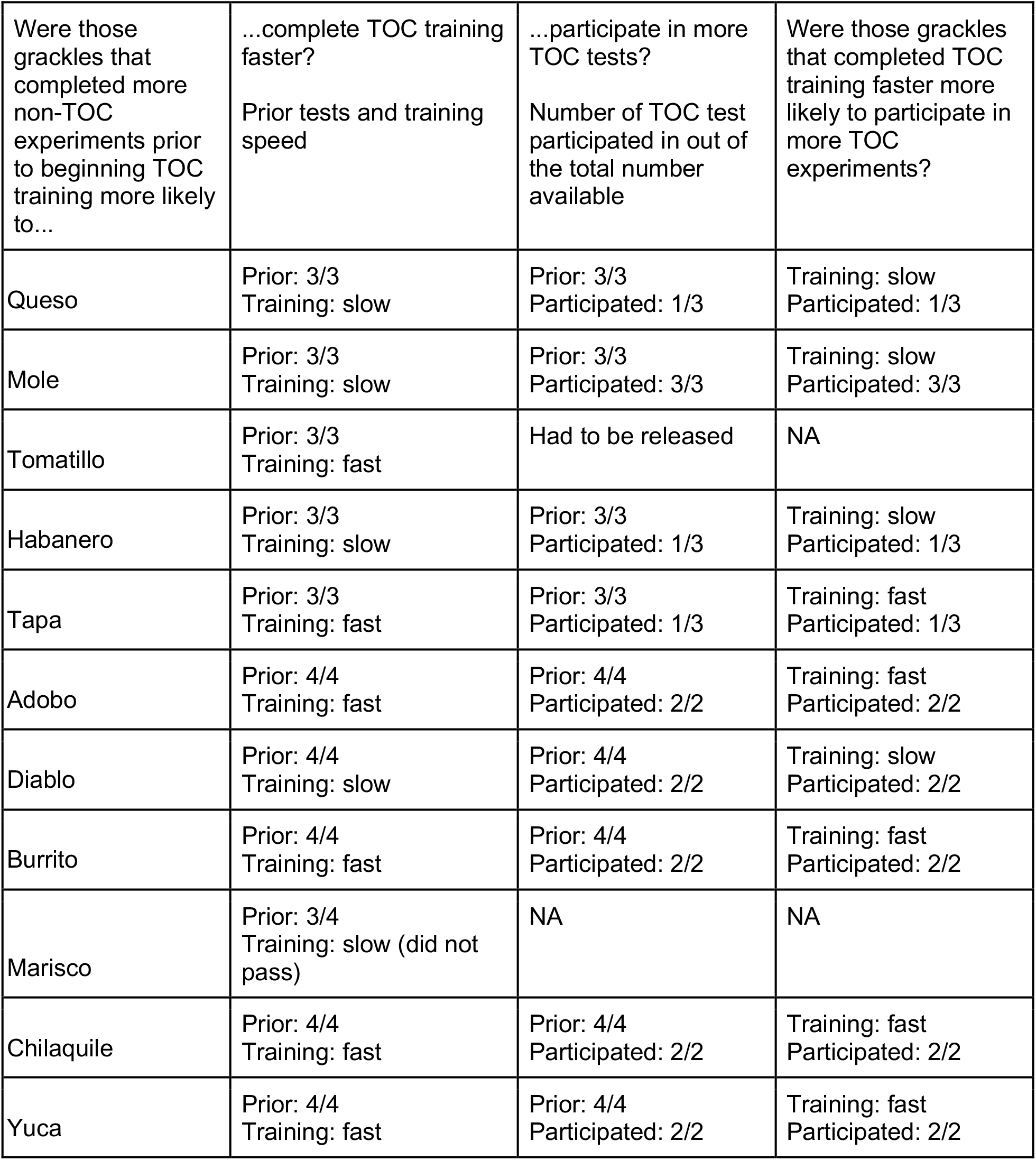
Summary data for the post-hoc questions (prior and participated x/y indicates that this individual participated in x out of the total possible y of the prior non-TOC experiments or TOC experiments). Only those birds who participated in enough TOC training to determine whether they were a fast (<13 days to pass training) or slow learner (13+ days to pass training) were included (n=11).

Question 2: Were those grackles that completed more experiments prior to beginning TOC training more likely to **participate in more TOC tests**? Answer 2: The number of previously completed experiments (three or four) was likely not related to the number of TOC experiments they completed (Table 2). The three grackles (Queso, Habanero, and Tapa) who did not complete all three TOC experiments did complete the maximum number of non-TOC experiments before beginning TOC training. The exception, again, was Marisco who completed only three of the four experiments attempted and did not participate in any TOC experiments because he never passed TOC training. Thus, the amount of prior experimental participation by individuals seemed unrelated to their ability to complete TOC experiments (i.e., feasibility of conducting TOC experiments given individual competence in prior experiments).

Question 3: Were those grackles that completed TOC training faster more likely to **participate in more TOC experiments**? The amount of training time might be predicted to inversely relate to the number of TOC experiments they complete, potentially because those individuals who require less training time may be more motivated to interact with and learn about the TOC, which could then also apply to the experiments involving the TOC. If such a relationship exists, this would be useful information for researchers who use TOCs because it would help them determine which individuals are more likely to complete testing, and thus which individuals to focus testing efforts on, if the research program is under time constraints. Answer 3: The duration of TOC training seemed unrelated to the number of TOC experiments grackles completed (Table 2). For example, Mole and Habanero took a long time to pass TOC training, but Mole completed all three TOC experiments, while Habanero completed only one. In contrast, Tapa was a relatively fast TOC learner, but only completed one out of three TOC experiments.

In summary, we do not appear to be able to use the number of previously completed tests as an indicator of likely TOC training speed or the likelihood of completing more TOC experiments, and TOC training speed was not an indicator of how many TOC experiments they might complete. Though we did not find any shortcuts for predicting grackle participation in TOC experiments, we offer these insights in the hopes that this information will help direct future efforts in productive ways.

## Programming TOC Experiments Using PsychoPy Programming Software

One of the largest barriers that hinders researchers interested in comparative cognition from conducting experiments on TOCs is that TOCs require tasks that must be written in some sort of computer programming language. There are a host of different options, many of which are not entirely user-friendly or intuitive for those without programming experience. Languages and programs that market themselves as more user-friendly, often require the purchase of costly licenses (e.g., E-Prime and MATLAB). To program tasks for our experiments investigating behavior in grackles (https://github.com/corinalogan/grackles/blob/master/README.md), and to study similar principles in pigeons and humans (https://pigeonrat.psych.ucla.edu/), we have had great success using PsychoPy and we highly recommend this software for those interested in building their own TOCs to study comparative cognition and beyond.

PsychoPy is a free and open-source application for programming experiments in the Python language and is compatible with most major operating systems (e.g., Windows, OS X, Linux) (Peirce, 2007; 2009; Peirce et al., 2019). Programs can be written directly in Python, or in the ‘Builder’ view, a graphical user interface (GUI) which allows for a simpler production of a wide variety of stimuli. The builder mode makes generating stimuli and documenting appropriate responses simple and also allows for ‘snippets’ of code to be added into the ‘Routine’ which allows for tremendous flexibility in terms of what can be presented and recorded during a task. There is also an active online support community (https://discourse.psychopy.org/) where hundreds of questions have already been addressed and new questions are swiftly attended to. Additionally, the latest version of PsychoPy (version 3 at the time of this publication) allows studies to be conducted online, which could be useful for studies where data are collected in different locations, for automatic online backup of the programs and data, and for easy sharing of programs among researchers (Bridges, Pitiot, MacAskill, & Peirce, 2020). There are additional advantages for researchers interested in non-human versus human comparisons on identical or slightly modified tasks. While it was outside the scope of this article to fully explain the capabilities of PsychoPy, we highly recommend that those interested consult a recently published textbook (Peirce & MacAskill, 2018), which details how to use the program for creating experiments.

Instead, we will focus on how to use PsychoPy to create tasks that involve the operations of external hardware to deliver outcomes (e.g., food delivery via the food hopper in our grackle TOC) which is typical of a TOC setup. We then provide basic templates and advice on creating programs specifically for studying comparative cognition—this should prevent some researchers from having to “reinvent the wheel.” For more information and examples of our code, including a sample Go/No-Go inhibition task, please consult the following page (https://github.com/corinalogan/grackles/tree/master/Files/TouchscreenPsychoPy2code).

### Setup: connecting the TOC to a food hopper

The first and perhaps most complicated step to setting up a TOC is establishing a connection between the program and an external hardware system that delivers some sort of outcome to the animal (e.g., food or shock). In our tasks, we used access to a food hopper as reinforcement. The food hopper was attached to a robotic arm (Pololu Maestro 12 Channel USB Servo, Pololu Robotics, Las Vegas, NV) that could be raised or lowered. Thus, in the very beginning of our program for a given experiment, a “snippet” of code was inserted to establish a connection between the program and the hopper. Given the universality of Python as a programming language, there may already exist online resources that explain how to control the external hardware using Python. That was the case for our Pololu system (https://github.com/FRC4564/Maestro). All experiments we conducted using this setup began with a brief 3 s routine that established a connection between the program and the hardware and then another 3 s routine that ensured the hopper was in the resting position out of reach from the subject (see Figure 2 for a snapshot of the experimental arrangement in PsychoPy).

**Figure 2.**
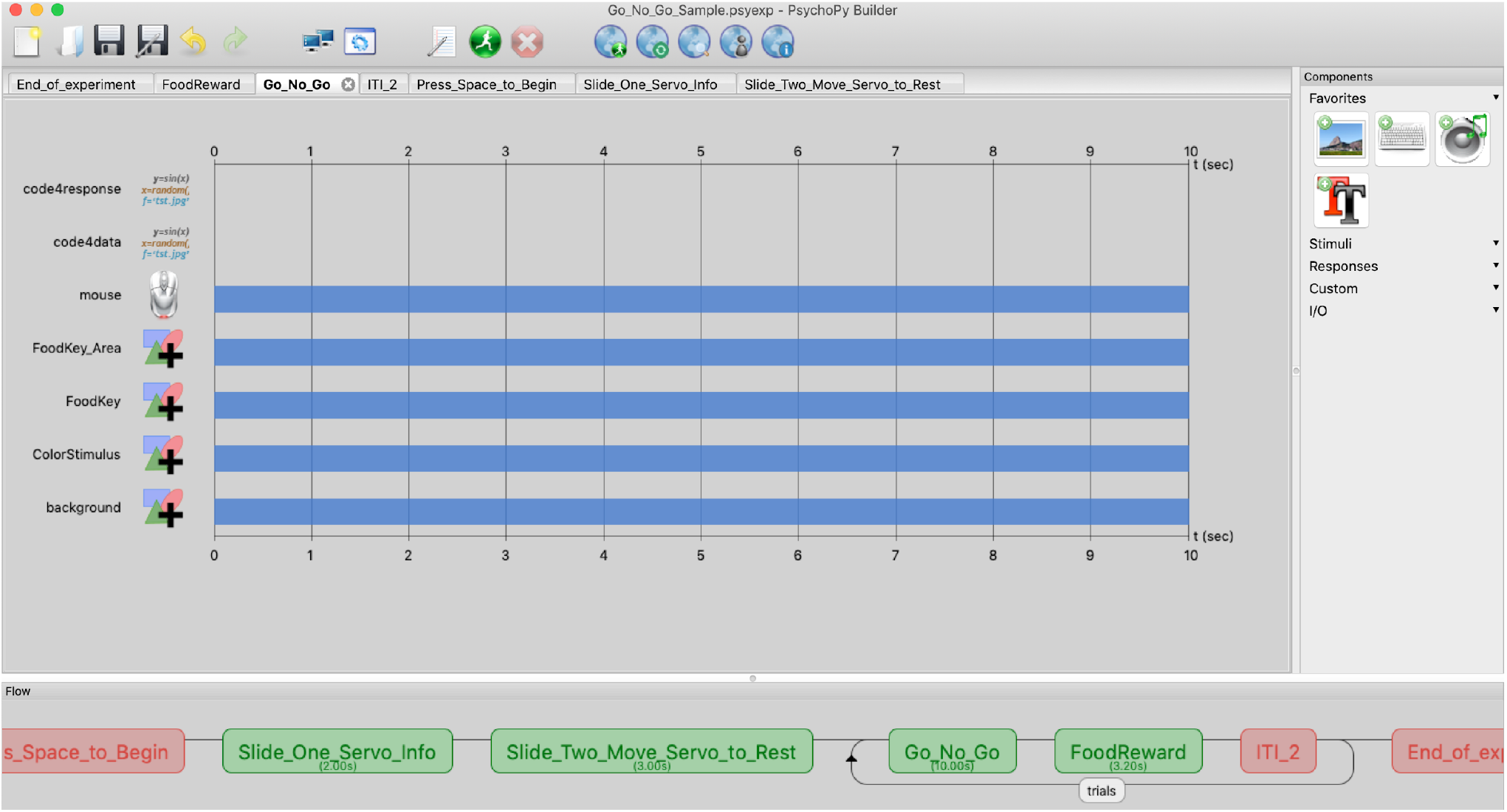
Basic experimental setup in PsychoPy. The “flow” (bottom bar) of the experiment represents the basic structure. The Press_Space_to_Begin “routine” allows time for the experimenter to initiate the experiment on command. The first two routines (Slide_One_Servo_Info and Slide_Two_Move_Servo_to_Rest) include code to establish a connection with the food hopper and then move it into the rest position. The rest of the experiment is generated within the loop titled, ‘trials’ (bottom bar).

### Basic Programming Outline

Designing a ‘Routine’ in PsychoPy builder mode is similar to creating a film in a video editing software in that it displays the temporal sequence of stimuli and responses that can occur throughout a given trial (see figure 2 as an example). Different stimuli can be presented during each routine and a number of different responses can be made. Additionally, the same routine can be used to create an infinite number of different trials because the routine can be conditionalized to an Excel file which determines which stimuli and responses will be presented (for more resources consult Peirce et al., 2019 & Peirce & MacAskill, 2018). We created a heavily annotated Go/No-Go sample program (https://github.com/corinalogan/grackles/tree/master/Files/TouchscreenPsychoPy2code/GoNoGo/2020-03AnnotatedGoNoGoCode) to illustrate these features. In that program, we created a routine whereby a discriminative stimulus was presented above a food key, and the correct decision to peck or not peck the food key led to a reward. The features (e.g., color) of the discriminative stimulus and whether or not it would be rewarded was entirely contingent on the elements within the Excel file (see Figures 3 and 4).

**Figure 3.**
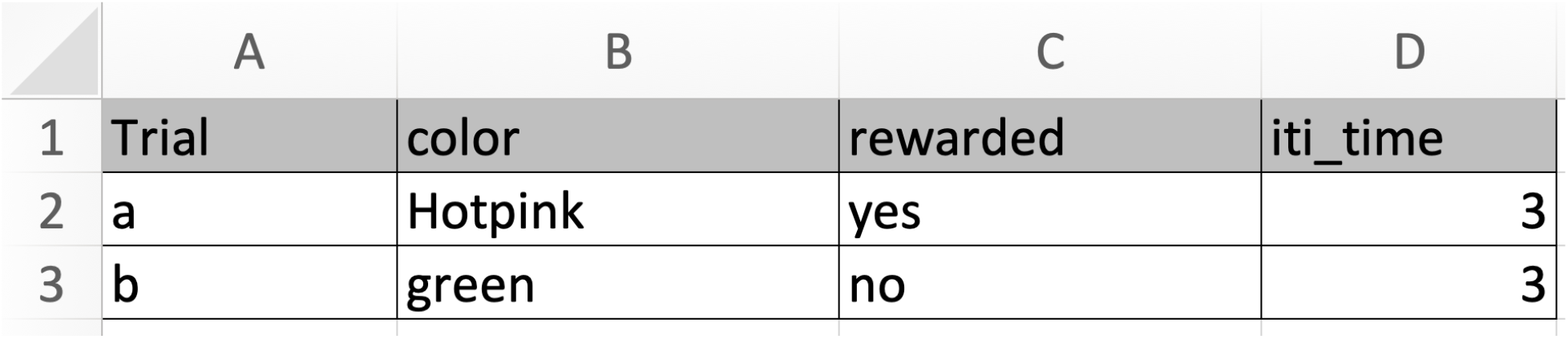
Setting up the conditions in an Excel file that is attached to the PsychoPy program as it runs. Psychopy will select a row from the attached excel file, and use the elements within that row to generate the trial. Thus, if row 2 were selected, the discriminative stimulus would be hot pink, a peck to the food key would be rewarded, and the inter-trial interval would be 3 seconds.

**Figure 4:**
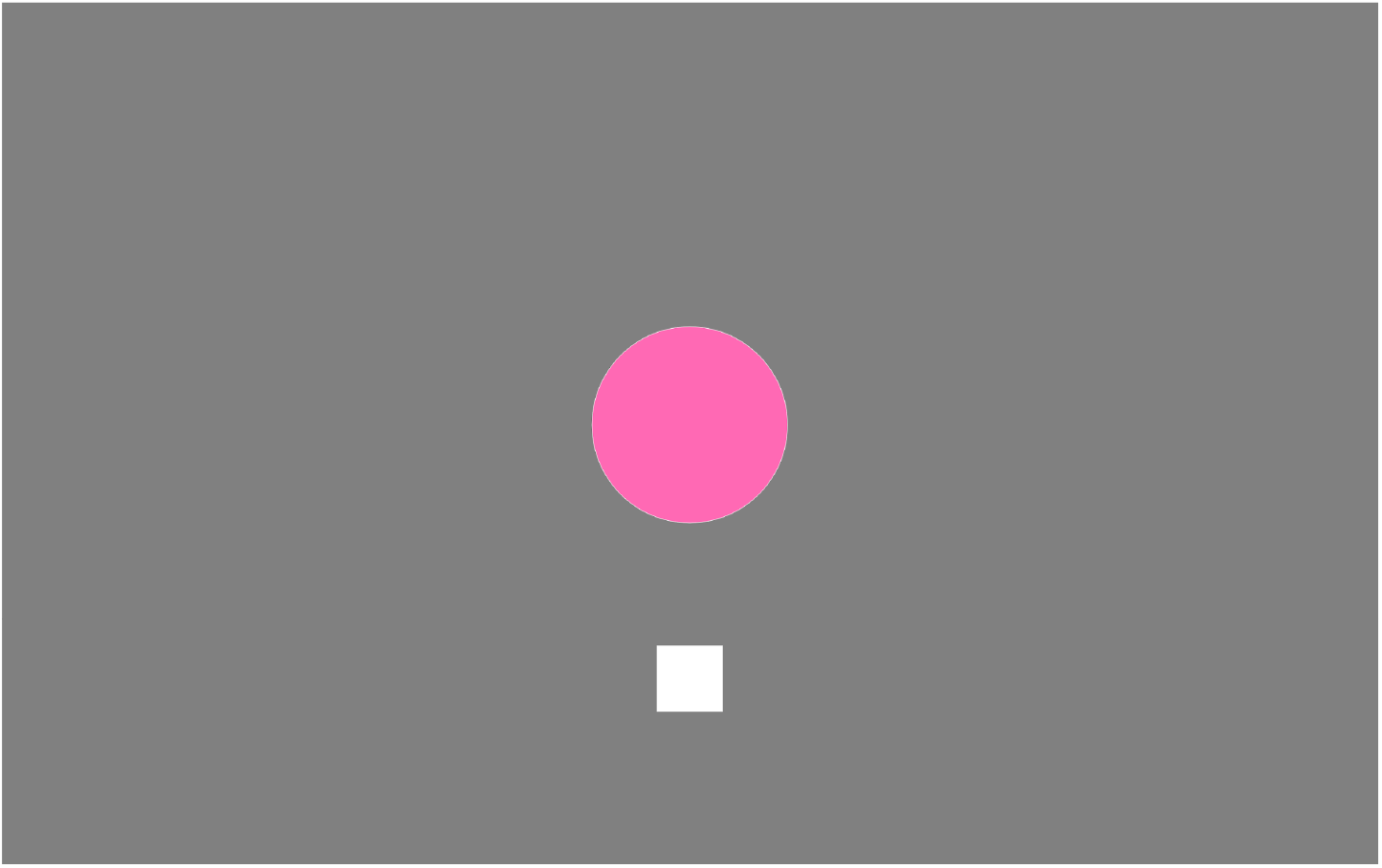
Example trial in PsychoPy. The color of the discriminative stimulus (circle) is dictated by the row chosen in the excel file, as is what happens when the food key is pecked.

Thus, for the majority of our experiments we created two main routines, one that included all of the desired features of a trial and the other that controlled the hopper and resulted in a food reward or non-reward depending on the response that was made. These two routines were enclosed in a loop that allowed us to determine the exact number of trials the animal would receive and which stimulus they would experience during those trials. Figure 2 provides an example of what this looked like in builder view, and an example of the code is provided in the Supplementary Material (S2). In the described sample Go/No-Go inhibition task, on some trials, a response to the food key terminated the routine and led to the next routine which was the food reward. Although PsychoPy is primarily used in an entirely graphical mode, extra capabilities can be achieved by specifying short snippets of custom code, written in the Python language, that run at specific times (such as at the beginning or end of the routine, or on every screen refresh). For example, if a response was made to the food key while the non-reinforced stimulus was presented, the trial ended and a new argument was created that we coded as “skip_food_reward = True”. The following food reward routine contained a statement to immediately terminate the trial and not offer a food reward if skip_food_reward = True (see S3). In other words, the pecking response on a No-Go trial initiated a clause that skipped the food reward routine that normally followed a trial. This resulted in no delivery of reinforcement as a result of the incorrect response. These simple snippets of code are intuitive even to those with little programming experience and are made easier to implement using the GUI.

### Food Reward / External Communication Routine

One of the most useful techniques involved simple stimuli provided by PsychoPy as proxies or representations for the movement of our external hardware (i.e., food hopper). Whenever we wanted to represent when the food hopper would be in the feeding position, we had a textbox appear on screen that said, “feeder is open”. When we wanted to represent the feeder being lowered to the resting position, a different text box read, “feeder is closing”. We then added a snippet of code that in essence posited: if the “*feeder is open”* textbox is on the screen, move the servo to the feeding position and if the “*feeder is closing”* textbox is on, move the feeder to rest. Thus, changing the duration of the food reward was as simple as changing the duration of the “feeder is open” textbox. We could also temporarily deactivate the code for the food hopper by adding a “#” in front of the argument (see S4). This allowed us to code experiments from any personal laptop (despite these computers not being connected to the hopper) anywhere in the world. Disconnecting the hopper also allowed the remote coder to test the program on the TOC without engaging the hopper. We could easily share programs among team members until the final program was agreed upon, then load it onto the TOC with the #’s removed and the text made invisible (by setting text font size to 0).

### Drawbacks of PsychoPy in Studies of Comparative Cognition

One noticeable drawback to using PsychoPy to operate TOCs and study comparative cognition is that each TOC must be operated by an individual computer. That is, while some commonly used software (e.g., MedPC by Med Associates, Inc.) allows for a single computer to operate multiple operant chambers, our setup requires that each screen is operated by its own computer. This may pose some practicality issues for those interested in running squads of animals on the same procedure, however it is less problematic for running only a handful of animals at a time. With that said, we are not aware of any technical reasons as to why PsychoPy cannot be used to control multiple monitors and external reinforcement machinery—but we are also not aware of any group who has successfully managed to construct such a setup.

## Concluding Remarks

Touchscreen-equipped operant chambers confer a number of advantages over typical operant chambers and traditional laboratory-based techniques. Despite the reviewed challenges, we expect TOCs to continue to grow in popularity in a number of scientific disciplines. As some traditional laboratory procedures appear to easily transfer to a TOC, while others do not, future research should explore the causes of these discrepancies. After several years of training wild-caught animals to operate TOCs, we have used this article to share the advantages and disadvantages of our various approaches, report data from our ongoing experiments, and provide advice and programming suggestions for researchers interested in similar pursuits. The programs used throughout this project (and data) are accessible on open-science platforms, and the move from traditional OCs and open-field procedures to TOCs represents another opportunity to make behavioral science more precise, transparent, and replicable.

## Acknowledgements

We thank Al Kamil and Debbie Kelly for brainstorming sessions on how to deal with the variety of problems we faced during training; Richard McElreath for project support; and our research assistants who helped trap the grackles and bring them into the aviaries: Aelin Mayer, Nancy Rodriguez, Brianna Thomas, Aldora Messinger, Elysia Mamola, Michael Guillen, Rita Barakat, Adriana Boderash, Olateju Ojekunle, August Sevchik, Justin Huynh, Jennifer Berens, Amanda Overholt, Michael Pickett, Mina Mohammed, Emily Blackwell, Kaylee Delcid, Brynna Hood, Samantha Bowser, Elise Lange, Sierra Planck, and Samuel Muñoz. For advice on using PsychoPy in this manuscript and technical support throughout this project, we thank Jon Peirce and Michael MacAskill. The research on wild-caught grackles was possible under the IACUC no. 17-1594R from Arizona State University, and through CL’s US Fish and Wildlife Service Scientific Collecting permit (number MB76700A-0,1,2), Bird Banding Permit from USGS (number 23872), and a Scientific Collecting permit from the Arizona Game and Fish Department (SP594338 [2017], SP606267 [2018], SP639866 [2019], and SP402153 [2020]). Benjamin Seitz is supported by National Science Foundation grant DGE-1650604. Aaron P. Blaisdell is supported by National Science Foundation research grant BCS-1844144.

## Supplementary Material

S1: Detailed Grackle Training Procedure, Tips, and Tricks

### The aviary set up for wild-caught grackles

The 12 adult grackles (Fajita, Mole, Tomatillo, Habanero, Chalupa, Tapa, Adobo, Diablo, Burrito, Marisco, and Yuca) and 2 juvenile grackles (Taco and Chilaquile; see Table 1) that underwent touchscreen training were caught in the wild in Tempe, Arizona, USA, from September 2018 through November 2019 and temporarily brought into aviaries for behavioral testing before being released back to the wild. All procedures were approved by University of Cambridge ethical review process. Each bird was measured, then color-marked with leg bands in unique combinations for identification, and blood samples were drawn prior to being placed in the aviary as part of other research (see Logan et al. 2019a, b; Blaisdell et al. 2019). Grackles were individually housed in an aviary (each 2.44 m long by 1.22 m wide by 2.13 m tall) at Arizona State University for a maximum of six months. All subjects had *ad libitum* access to water at all times and were fed Mazuri Small Bird maintenance diet *ad libitum* during non-testing hours (minimum 20 h per day), and other, more preferred food items (e.g., goldfish crackers) during testing. Individuals were given a minimum of three days to habituate to the aviaries and then their test battery usually began on the fourth day. Birds were usually tested six days per week, therefore if their fourth day in the aviaries occurred on a day off, then they began testing on the fifth day instead. After engaging in three to four experiments (experiment order was pseudorandomized such that some individuals received TOC training earlier than others), grackles were then habituated to and trained to use the TOC, and subsequently participated in one to three TOC experiments.

### The Touchscreen-equipped Operant Chamber training procedure

We trained grackles to interact with the TOC through three distinct steps: habituation to the TOC, hopper training, and training to peck the screen. Our PsychoPy code for the below procedures (as well as the TOC experiments) is available at https://github.com/corinalogan/grackles/tree/master/Files/TouchscreenPsychoPy2code.

**Table S1.** Outline of the three consecutive steps involved in training grackles to use a TOC.

| **1. Habituation to TOC apparatus** | **2. Hopper training** | **3. Touch the screen for food** |
| --- | --- | --- |
| TOC present, but not turned on | TOC present, food hopper is active, touchscreen is turned off | TOC present and all pieces (food hopper and touchscreen) are on |

#### 1. Habituation to TOC apparatus

Grackles were first habituated to the presence of the TOC. The TOC was placed in each aviary overnight with maintenance diet only accessible from the hopper (container with food that can automatically move up or down to grant or deny food access) and atop the platform in front of the hopper (i.e., food is only available on or in the TOC and no food is available from their regular food dish away from the TOC). This was done for between 0 - 12 nights, with the touchscreen being left in the aviary for additional habituation if grackles continued to avoid approaching it during daytime active training sessions. Grackles that did not receive the TOC in their aviaries overnight were either birds that the experimenters felt could destroy the apparatus when left alone with it (e.g., Adobo) or those that participated readily in daytime sessions without prior overnight habituation (e.g., Yuca and Chilaquile). We actively encouraged the birds to approach the TOC monitor and eat Goldfish crackers out of the raised hopper during daytime sessions by sprinkling reward food (i.e., crackers) on the TOC platform in front of the monitor and hopper. Once grackles were comfortable eating from the raised hopper, we progressed to hopper training (below) to habituate the grackles to the sound and movement of the hopper and to learn they have a limited time period in which to access the food when the hopper is raised.

#### 2. Hopper training

Grackles were trained to associate the sound and movement of the hopper being raised with the availability of food. To do so, we removed all food from the platform such that food (e.g., crushed up Goldfish crackers) was only available from within the hopper, and we began moving the hopper to deliver food to the birds when they approached it. This was made possible by remotely controlling the computer (*a DELL Inspiron 15*) attached to the TOC using a separate computer (*experimenter’s computer*) outside of the testing area (Figure 1). Specifically, we used a free trial version of TeamViewer software (version 15.2.2756; www.teamviewer.com) to remotely control and run all training and testing programs on the TOC computer via a wifi connection. A number of free remote operating systems exist (e.g., Google Chrome Remote Desktop) that would conceivably serve the same function. We created a PsychoPy program to move the food hopper up or down with every press of the space bar on the experimenter’s computer (“hopper training” program; file name: <u>1.Press_Space_for_food_2.Basic_mag_training_nostartscreen.psyexp</u>). We habituated grackles to the sound and movement of the hopper by first lowering the hopper remotely after an individual had eaten from the raised hopper. Once an individual was habituated to the sound and movement of the hopper lowering after reward delivery, the experimenter switched to raising the hopper remotely as the bird approached the hopper. When the bird learned that the noise of the hopper moving meant that food was available, and they consistently approached and ate from the raised hopper, they progressed to the next stage of training in which they learned to touch the screen for the food reward (below). It took grackles 3-25 days to pass hopper training (*n*=11, Table 1). We attempted hopper training with an additional three grackles, but two had to be released back to the wild before their training was complete and one did not pass hopper training (see “Hopper Training” in Table 1).

**Table S2.** Hopper training summary data. The number of days is the total number of days included in the period from the first to the last training day (i.e., it is not the number of days on which training occurred), the number of sessions and trials include those where the bird did not participate. Note that when a number has a +, this means that the detailed data for the initial trials/sessions were not recorded. This is because we did not realize right away that collecting this data at the trial level would be useful for understanding how to better train grackles on the apparatus.

| Bird ID | Passed training | Number of days | Number of sessions | Number of trials |
| --- | --- | --- | --- | --- |
| Fajita | No | 2 | 3 | 16 |
| Queso | Yes | 6 | 3+ | 25+ |
| Mole | Yes | 25 | 5+ | 21+ |
| Tomatillo | Yes | 5 | 4+ | 20+ |
| Habanero | Yes | 4 | 4+ | Not recorded |
| Chalupa | No | 5 | 6+ | 9+ |
| Tapa | Yes | 2 | 2 | 22 |
| Taco | Yes | 10 | 14 | 38 |
| Adobo | Yes | 4 | 6 | 81 |
| Diablo | Yes | 16 | 22 | 44+ |
| Burrito | Yes | 3 | 5 | 20+ |
| Marisco | No | 54 | 67 | 98 |
| Chilaquile | Yes | 3 | 4 | 40+ |
| Yuca | Yes | 6 | 12 | 56 |

#### 3. Touch the screen to obtain food

Grackles were then trained to touch the screen to gain access to a food reward. We created a program that presented a small (2 cm × 2 cm) digital white square on the screen (“white square training” program, file name: <u>4.Food_Key_Only_2FullControl.psyexp</u>). Pecks made directly to this stimulus resulted in an automatic 5 s of food access from the raised hopper, or the experimenter could press the spacebar on their laptop to raise the hopper manually. We initially intended to use a mixed Pavlovian and Instrumental autoshaping procedure to encourage birds to engage with the TOC to obtain food rewards (i.e., the birds could peck a stimulus to receive the reward [Instrumental] and a trial always ended in reward regardless of their behavior [Pavlovian/Autoshaping]), but it became immediately clear when we started training that this would not work with the grackles who were not interested in interacting with the touchscreen unless they were hand-shaped to do so. Instead, experimenters employed more basic hand-shaping procedures to encourage grackles to engage with the screen (i.e., the experimenter had to be present outside of the aviary to trigger the hopper to reward the bird for incrementally correct behaviors that eventually led to their touching the digital white square). Grackles were rewarded (by the experimenter cueing the hopper to move to the available position so the bird could eat) for first putting their bill on the screen near the digital white square, then closer to the square, until they were touching the square on their own and able to trigger the hopper themselves.

However, the hand-shaping technique of remotely triggering the food hopper to get the grackle to peck the digital white square was not enough to get them to successfully engage with the digital white square. All subjects required additional hand-shaping methods. One method for trying to get the bird to peck the digital white square on the screen was to have the experimenter demonstrate by touching the square with their finger, which would trigger the hopper, and then the experimenter could reset the hopper and show the correct behavior a few times. This drew the bird’s attention to the screen, often resulting in them coming to the screen to explore it (i.e., stimulus enhancement), thus increasing their chance to learn something about the stimulus. The primary method used to encourage engagement with the screen was to put a piece of clear tape that contained bits of crushed Goldfish crackers over the digital white square on the screen. This resulted in grackles grabbing at the crackers, and coincidentally touching the screen, which triggered the hopper to raise. However, due to the sensitivity of the TOC, the tape on the screen often disrupted the “white square training” PsychoPy program because the program registered the tape as a touch and locked the hopper in a raised or lowered position. A more successful method for training the grackles to interact with the screen involved using the “hopper training” PsychoPy program, which allowed experimenters to control the hopper and it did not have a digital white square on the screen. Combined with this program, experimenters taped a paper white square onto the screen and subsequently Goldfish crackers were taped on top of the paper square. When the grackle touched the Goldfish or paper white square, the experimenter remotely controlled the hopper to make the food accessible. Once grackles were consistently pecking at the crackers taped to the paper white square, we decreased the size of the cracker crumbs until the grackle learned it would receive a reward from the hopper for simply pecking the paper white square. After the bird consistently pecked the paper white square taped over the screen and ate from the hopper, we removed the paper white square and moved to the digital “white square” PsychoPy program. If the bird did not peck the digital white square, we continued to intersperse paper white square and digital white square trials until they did.

Although we were able to find an efficient method of training most grackles to use the TOC, throughout this discovery process there were many individual differences in the efficiency and consistency with which grackles learned to interact with the screen that required some flexibility in the training protocol. For example, some birds made pecks to the digital white square in such a way that they were not registered by the touchscreen, therefore it was important for the experimenter to remotely raise the hopper when the bird made an accurate peck to ensure the association was made between touching the white square and the food reward. Additionally, some birds pecked anywhere on the screen rather than specifically on the digital white square, so we reverted to taping food over the square to strengthen the association between the shape and the food. Some individuals took a long time to shape to the digital white square and required a significant amount of time to retrain associations before the grackle was able to consistently touch the digital white square to elicit a food reward. For example, Habanero’s later training sessions started with him pecking the paper white square taped to the screen because he didn’t touch the digital white square, even though he had touched the digital white square and triggered the hopper in previous sessions. Even during later training stages, other individuals who had been comfortably interacting with the TOC sometimes reverted to fearful behavior where they would not approach the TOC during a session, or they would approach quickly and jump back rather than remaining close to interact with the stimuli on several sequential trials. In these cases, we reverted to paper white square training or hopper training until they again appeared comfortable with the TOC.

A total of 11 grackles received white square training and 10 of them passed (one had to be released back to the wild before his training was complete; see White Square Training in Table 1). It took 2-43 days to complete this training, however one grackle required 58+ days because of a medical procedure he underwent in the middle of the training period.

**Table S3.**
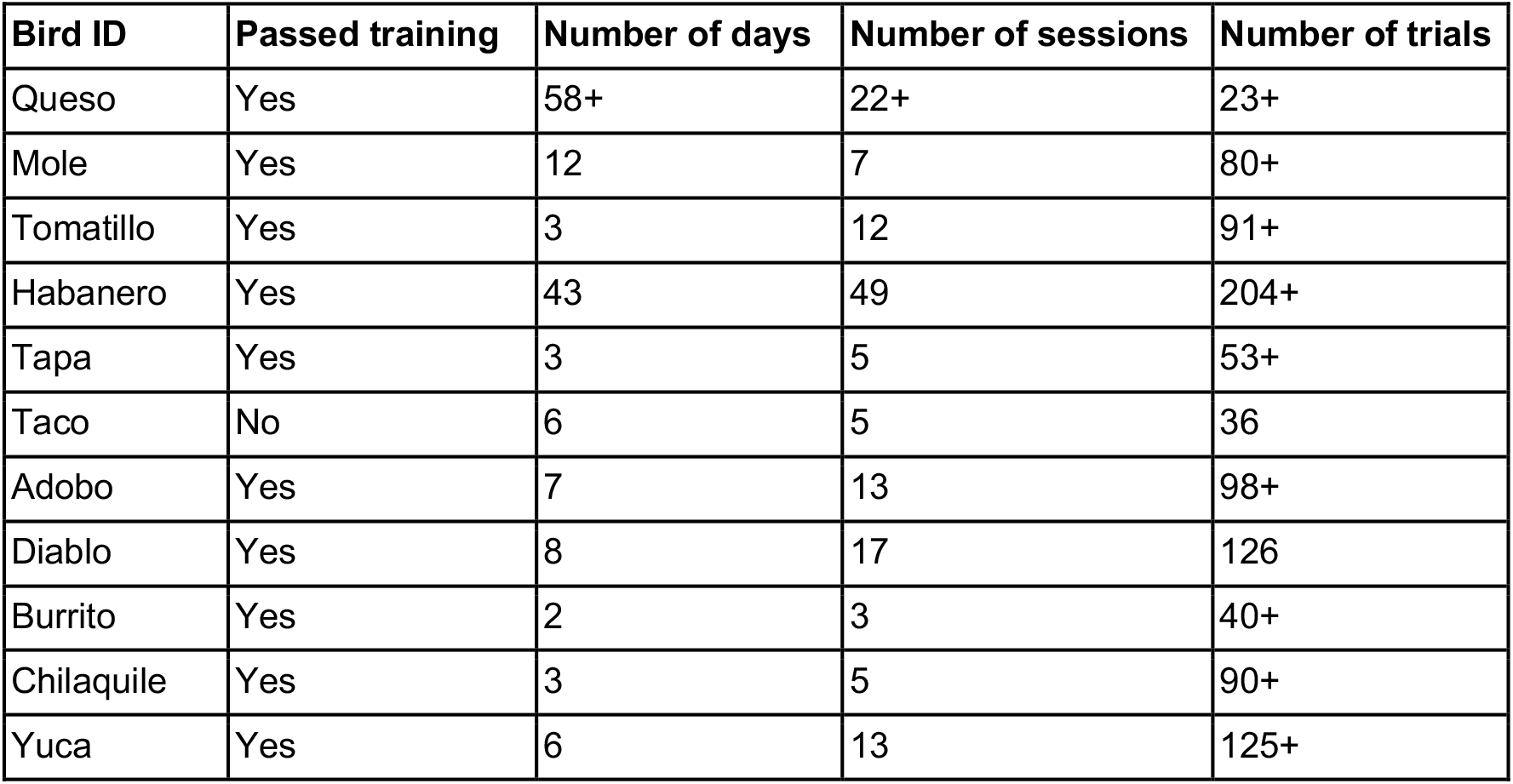
White square training summary data. The number of days is the total number of days included in the period from the first to the last training day (i.e., it is not the number of days on which training occurred), the number of sessions and trials include those where the bird did not participate. Note that when a number has a +, this means that the detailed data for the initial trials/sessions were not recorded. This is because we did not realize right away that collecting this data at the trial level would be useful for understanding how to better train grackles on the apparatus.

#### Additional training programs that were discontinued

We originally thought that it might be more ecologically relevant for the grackles to learn to interact with the TOC by learning to peck a shape that was moving around on the screen, simulating a moving insect (grackles often catch insects on the fly). Therefore, we made the “Moving target training” (file name: <u>moving_stim_.psyexp</u>) program and tried it on two of the first grackles to be trained. Grackles were not more likely to peck a moving white circle without the extra hand-shaping training described above. Therefore, we continued training only with the white square training program because it was much easier to implement the training methods above if the stimulus did not change positions on the screen (e.g., placing paper and crackers over the location of the digital white square).

For some of the TOC experiments, we were planning to have a trial start key. The function of the trial start key was that a trial would only start when a bird was paying attention and ready to participate, and the bird could indicate this by pecking the trial start key to initiate each trial. When the trial start key was pecked, the test trial began. Two grackles were given (and passed) trial start key training (file name: <u>5. Trial_Start Key Shape.psyexp</u>). However, as these birds moved through the experiments, the trial start key was an extra (unrewarded by food) step before the start of the trial and it negatively impacted their motivation to participate. To avoid these problems with motivation, we decided it was not necessary to include the trial start key because the experimenter could determine when the bird was attending to the screen and trigger the trial to start from the aisle of the aviary using their computer. We then removed the trial start key from all TOC experiments. While removing the start key also removes part of the automation of this experiment, which is a large benefit of a TOC, we do not think that grackles would be able to complete experiments on TOCs in a completely automated way. This is largely due to the motivational issues that we have encountered, although this may be confounded by our open setup, instead of a smaller enclosure which could theoretically facilitate engagement. Thus, for our experiments with grackles, the ability to remotely initiate trials was a great benefit to the TOC approach. Many researchers working with wild-caught individuals, or implementing TOC experiments in the wild will likely have similar problems. Therefore, we included exhaustive details of our experience to showcase the flexibility of TOC methods and to facilitate other research with similar subjects.

Once grackles had passed white square training, they were ready to begin participating in experiments with the TOC. However, we encountered a few additional issues. Originally, our reversal learning TOC experiment included two different colors and required color discrimination. However, the first two birds that passed TOC training and moved on to the reversal TOC experiment exhibited avoidance and/or fearful behaviors when presented with a light purple circle and a dark purple circle simultaneously on the screen. They refused to come back to the TOC area even though we tried to habituate them to a variety of colors and shapes on the screen. Therefore, we changed the reversal TOC experiment to instead discriminate between two different white shapes.

**S2:**
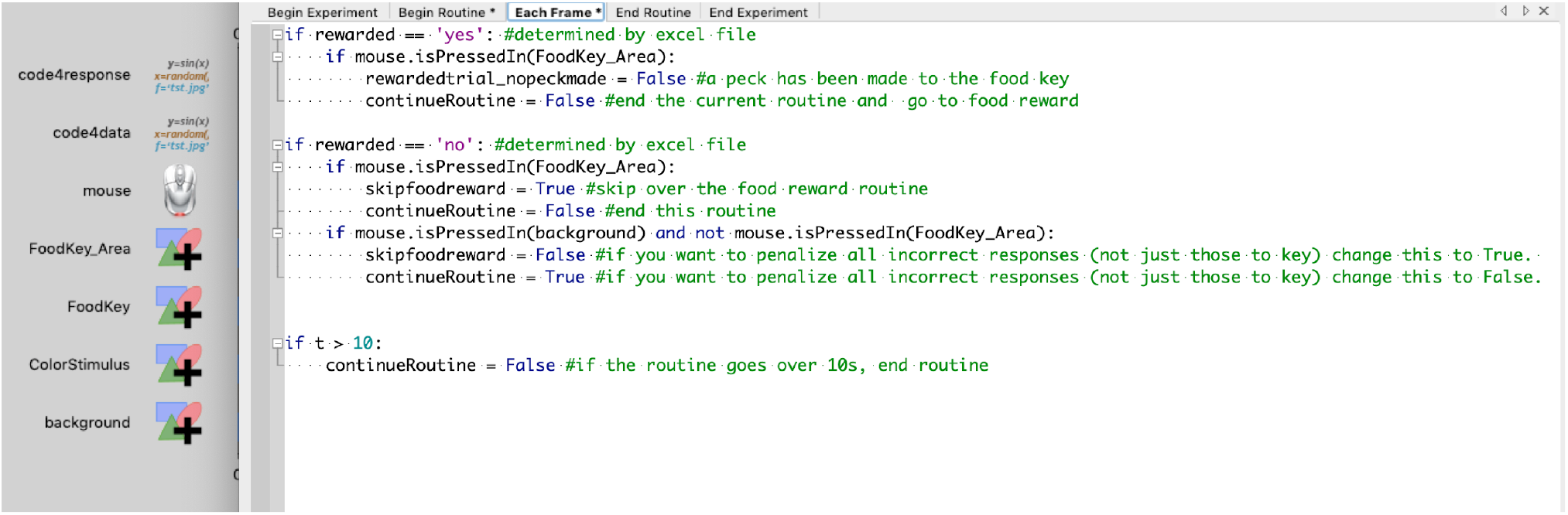
Screenshot of Go/No-Go code in PsychoPy. This code governs the Go/No-go trial and determines whether the subsequent foodreward routine should continue or be skipped over.

**S3:**
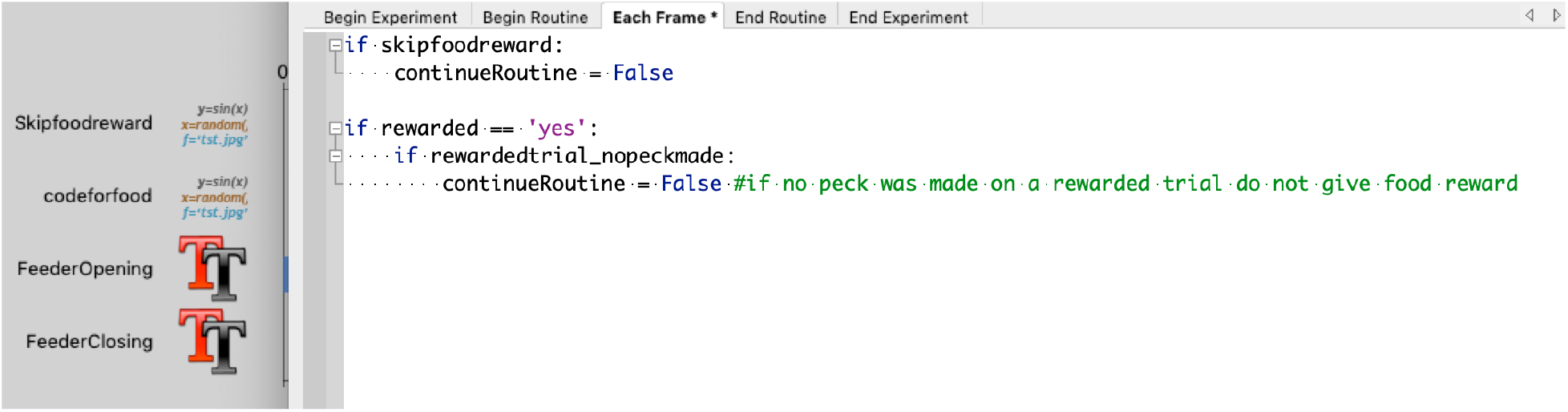
Screenshot of the food reward code in PsychoPy. If the cause skipfoodreward has been made true, the routine immediately ends and no food is delivered.

**S4:**
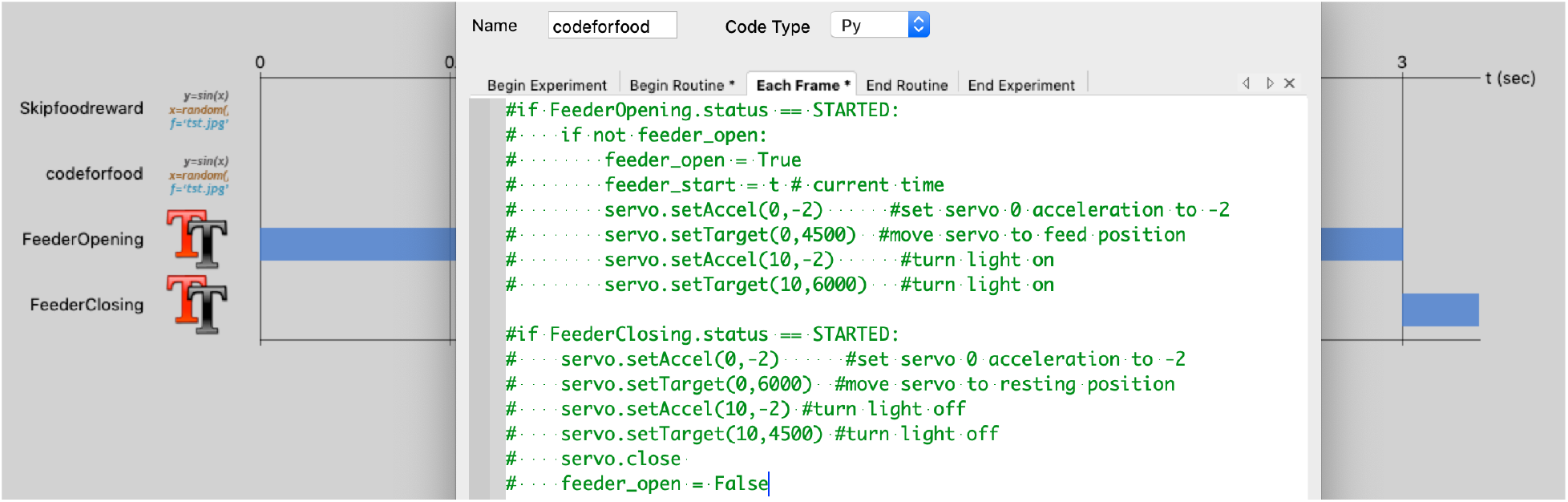
Screenshot of code to move the robotic food hopper (also called Servo) in PsychoPy. By making the hopper movements contingent on more simple elements like a textbox, we were able to test the program on computers not hooked up to the actual TOC. To increase the duration of the food reward, we could simply adjust the duration of the FeederOpening text element as well as the start time of the FeederClosing text element.

## Notes

### Competing Interest Statement

The authors have declared no competing interest.

